# Syntrophic H_2_ production enhances the performance of primarily acetate-supplemented reactors treating sulphate contaminated solutions

**DOI:** 10.1101/2021.05.12.443249

**Authors:** Tomas Hessler, Susan T.L. Harrison, Jillian F. Banfield, Robert J. Huddy

## Abstract

Biological sulfate reduction (BSR) represents a promising bioremediation strategy, yet the impact of metabolic interactions on performance has been largely unexplored. Here, genome-resolved metagenomics was used to characterise 17 microbial communities associated with reactors operated with defined sulfate-contaminated solutions. Pairs of reactors were supplemented with lactate or with acetate plus a small amount of fermentable substrate. At least thirty draft quality genomes, representing all the abundant bacteria, were recovered from each metagenome. All of the 22 SRB genomes encode genes for H_2_ consumption. And of the total 163 genomes recovered, 130 encode 321 NiFe and FeFe hydrogenases. The lactate-supplemented packed-bed bioreactor was particularly interesting as it resulted in stratified microbial communities that were distinct in their predominant metabolisms. Pathways for fermentation of lactate and hydrogen production were enriched towards the inlet whereas increased autotrophy and acetate-oxidizing SRB were evident towards the end of the flow path. We hypothesized that high sulfate removal towards the end of the flow path, despite acetate being an electron donor that typically sustains low SRB growth rates, was stimulated by H_2_ consumption. This hypothesis was supported by sustained performance of the predominantly acetate-supplemented stirred-tank reactor, which was dominated by diverse fermentative, hydrogen-evolving bacteria and low-abundance SRB capable of acetate and hydrogen consumption. We conclude that the performance of BSR reactors supplemented with inexpensive acetate can be improved by the addition of a low concentration of fermentable material due to stimulation of syntrophic relationships among hydrogen-producing non-SRB and dual hydrogen- and acetate-utilising SRB.

## Introduction

Sulfate contaminated wastewater is generated by a number of industries but posing the greatest environmental threat is acid rock drainage (ARD) arising from both current and historic mining activities. Abiotic and microbial oxidation of sulfidic ore produces low pH solutions with elevated sulfate and heavy metal concentrations (Silverman and Ehrlich, 1964). Although physical and chemical treatments are effective at remediating major ARD sites (e.g.mine water discharged from active mine workings), these processes are not suitable for the treatment of very common localized low-flow ARD because they demand substantial infrastructure and have high operating costs.

Decades of study have shown BSR to be a promising low-cost approach for the bioremediation of diffuse sources of lower volume and less acidic ARD (Kaksonen and Puhakka, 2007; Zagury and Neculita, 2007). BSR is catalysed by a diverse group of anaerobic microorganisms known as sulfate-reducing bacteria (SRB). Sulfate present in the ARD is used by the SRB within bioreactors as a terminal electron acceptor coupled to the oxidation of a supplied electron donor, resulting in the generation of sulfide and bicarbonate (Muyzer and Stams, 2008). The generated sulfide can be used to precipitate heavy metals in solution or be partially oxidized to elemental sulphur (van Hille et al., 2016), a value-added product. The choice of supplied electron donor largely dictates the cost of an ARD-remediating BSR process (Papirio et al., 2013). Lactate and ethanol are often studied due to the relatively high SRB growth rates these support and several operating conditions required to favor the growth of SRB over fermentative microorganisms have been optimised. A major limitation of BSR processes operated with various volatile fatty acids (VFAs) and complex carbon sources is the frequent build-up of acetate in the effluent of these reactors, even in the presence of high sulfate concentrations (Omil et al., 1997; Lens et al., 2002). The difficulty to establish an active acetate-utilising SRB community in these reactor systems has puzzled researchers for some time (Oude Elferink et al., 1995). Acetate is of particular interest as an electron donor for sustainable ARD remediating BSR processes due to its low cost compared to other electron donors and its potential to be sourced from various waste streams. Therefore, understanding the factors which enhance acetate-utilising SRB’s growth within BSR reactors is important for the success of acetate-supplemented BSR systems as well as the efficient utilisation of other electron donors.

Despite the promise, few ARD-remediating BSR processes have been implemented and operated at scale. The successful operation of BSR reactors faces several challenges, including the efficient utilisation of the supplied electron donor due to competition between SRB and other microorganisms within the microbial community. Further, achieving high reaction rates by overcoming the relatively low growth rates of SRB has been difficult (Harrison et al., 2014). These challenges are intimately linked to the microbial community dynamics and the metabolic potential of the organisms in these systems, both of which remain little studied.

SRB have been extensively investigated in natural environments (Jørgensen, 1982; Bowles et al., 2014), as these microorganisms play important roles in biogeochemical cycling. Pure and mixed SRB cultures have undergone thorough kinetic growth studies (O’Flaherty et al., 1998; Moosa et al., 2002; Oyekola et al., 2010) that have enabled the development of mathematical models used to optimise BSR processes (Cassidy et al., 2015). Complementing these culture-based studies, marker gene (e.g., the 16S rRNA gene) tracking has been used to uncover the factors that impact the microbial community composition within BSR reactor systems (Icgen and Harrison, 2006; Dar et al., 2008; Nielsen et al., 2018). However, recent genome-resolved metagenomic surveys of subsurface sediments and aquifers have revealed that SRB are more diverse than previously thought (Anantharaman et al., 2018). This raises questions around the suitability of using only 16S rRNA gene surveys for the identification of SRB within bioreactor communities.

Here, we report the genome-resolved metagenomic characterization of microbial communities associated with six continuously operated BSR reactors supplemented with lactate and with acetate as electron donors. These bioreactors were intended to follow neutralisation and heavy metal removal steps in the ARD remediation process and were, therefore, operated with neutral, defined drainage. We employed up-flow anaerobic packed bed reactors (UAPBR), linear flow channel reactors (LFCR) and continuous stirred-tank reactors (CSTR). UAPBRs are plug-flow governed configurations commonly used in BSR research (Kaksonen and Puhakka, 2007). These reactors enable high biomass retention in the form of biofilms that colonise the packing material. Plug-flow results in stratification of the nature and magnitude of the reactions along the reactor length. The LFCR is a newly-developed reactor configuration that achieves complete mixing passively within a single hydraulic retention time (HRT; Marais et al., 2020). Carbon microfibers were incorporated into this configuration to support biofilm formation. In contrast, CSTRs are well-mixed reactors with low surface area to volume ratios which result in predominantly planktonic microbial communities. Within this study, the CSTR systems allow the investigation of planktonic microbial communities in the absence of biofilm communities, and thereby rapid community dynamics in response to operating conditions, governed dominantly by biokinetics. The distinct selective pressures which resulted from the differences in reactor hydrodynamics, supplied electron donors and their capacity to support microbial biofilms was intended to lead to the divergence of the composition of these microbial communities from each other and the original inoculum.

## Results and discussion

### Bioreactor operation and sampling

Duplicate UAPBRs, LFCRs and CSTRs, each supplemented with acetate and lactate separately, were inoculated with a diverse sulfidogenic microbial culture. The reactors were operated continuously at a four-day hydraulic retention time (HRT). Total genomic DNA from the planktonic and biofilm microbial communities from these bioreactors was sampled, in duplicate, after the onset of steady-state conditions, between day 463 and 514 of continuous operation. The reactors displayed various degrees of sulfate conversion and degradation of VFAs. The observed sulfate conversion was lowest in the solely planktonic cell supporting, acetate-supplemented, CSTR (45%) and highest in the lactate- and acetate-supplemented UAPBRs (96 and 99%, respectively). The observed sulfate reduction was putatively linked to the oxidation of various quantified VFAs. Some of these reactor performance data have been previously reported (Hessler et al., 2018, 2020) and can be found in the supplementary data (Table S1). Duplicate samples were taken from the acetate- and the lactate-supplemented CSTRs for metagenomic analysis. Only one sample type was collected. as these represented well-mixed and completely uniform environments that support a primarily planktonic microbial community. Replicates of the planktonic and biofilm communities were sampled from the well-mixed LFCRs. The planktonic cell communities from the plug-low governed UAPBRs were sampled from each of the three sequential zones of the reactors, and matrix attached biofilms were sampled from the inlet- and effluent zones.

### Genome recovery

The 34 metagenomes (17 samples in duplicate), representing the initial inoculum and reactor microbial communities, were individually assembled and 163 draft microbial genomes were reconstructed. Based on the inventory of bacterial and archaeal single copy genes, the average estimated completeness of the recovered genomes was 95%, with 127 genomes having an estimated completeness of >90.0% and <5.0% contamination. The average and minimum proportion of the reads from each metagenome that mapped to binned contigs (≥1000 bp) was 89 and 85%, respectively. Thus, the genomes well represent the organisms present in the bioreactors. The bioreactor communities are relatively simple in composition, with an average Shannon index of 3.0. However, many of the genomes were for organisms found to be dominant in only a subset of the samples. We attribute the large number of high-quality recovered genomes to the heterogeneity in the physiochemical conditions generated across the bioreactor environments.

The reactors were supplemented with the methanogenic inhibitor BESA at the time of inoculation to provide an initial advantage to acetate-oxidising SRB over any methanogens for available acetate. Thus, although a single archaeal genome was recovered in the inoculum, archaea were almost completely undetectable in the reactor communities. The remaining 162 bacterial genomes were classified to 16 phyla but were predominantly assigned to Alphaproteobacteria (14 genomes), Betaproteobacteria (6), Deltaproteobacteria (22), Gammaproteobacteria (12), Epsilonbacteria (5), Firmicutes (29), Bacteroidetes (28), Synergistetes (15) and Spirochaetes (12) (Figure 1).

**Figure 1.**
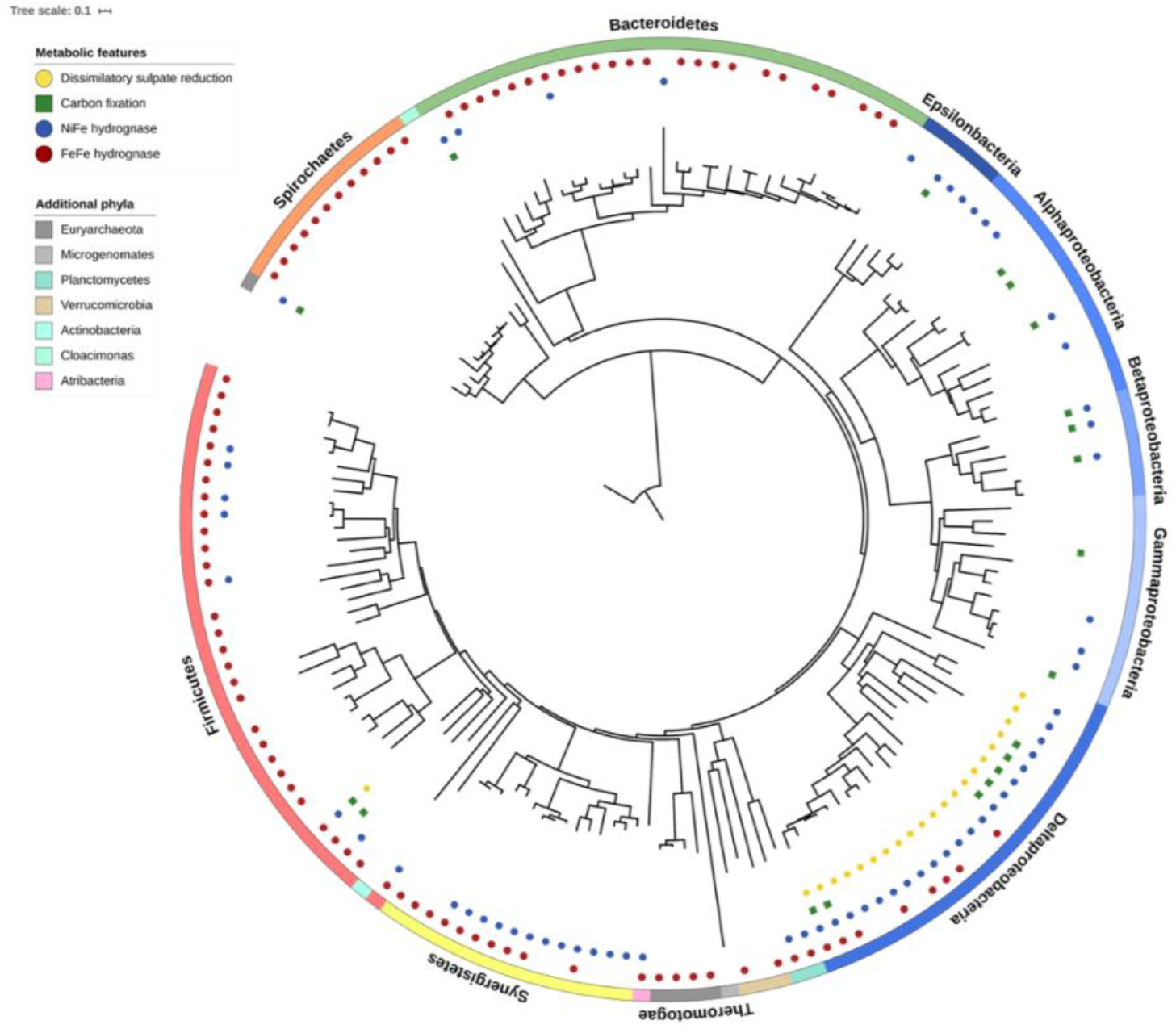
Phylogenetic tree of the 163 microbial genomes recovered from the inoculum and bioreactors of this study. The phylogenetic tree was constructed based on the concatenated alignments of 16 ribosomal proteins. The phyla to which each lineage belongs is annotated. Coloured symbols represent genome-inferred metabolic features. All identified SRB encoded NiFe hydrogenase (predominantly group 1b NiFe hydrogenases associated with H_2_ uptake) and several SRB genomes encoded the autotrophic Wood-Ljungdahl pathway. Many other Proteobacteria and Synergistetes encoded NiFe hydrogenase whilst FeFe hydrogenases were widespread throughout the remaining genomes recovered from these bioreactors.

### Microbial replication rates across reactor environments

Indices of replication (iRep) were determined to evaluate microbial replication rates across the different reactor environments (Figure 2). Most of these reactor communities showed narrow and similar distributions in represented iRep values. The narrow distributions indicate that the microbial communities were relatively stable at the time of sampling. Where biofilms were present, the median iRep values for bacteria in biofilms were generally lower than those of bacteria in the planktonic communities that occupied the same reactor zones, although the differences were not statistically significant. The generally comparable replication rates for planktonic and biofilm-associated bacteria is of particular interest as biomass quantification found biofilms contribute up to 100-fold more cells per reactor volume than planktonic communities within the same reactor zone (Figure S2). In addition, SRB were, on average, twice as abundant in biofilms compared with planktonic communities. The contribution of these biofilms to biomass retention within the reactors, the representation of SRB in these biofilms, and the similar iRep distributions between planktonic and biofilm communities demonstrates that the biofilms within these reactors likely contributed substantially to the observed sulfate reduction.

**Figure 2.**
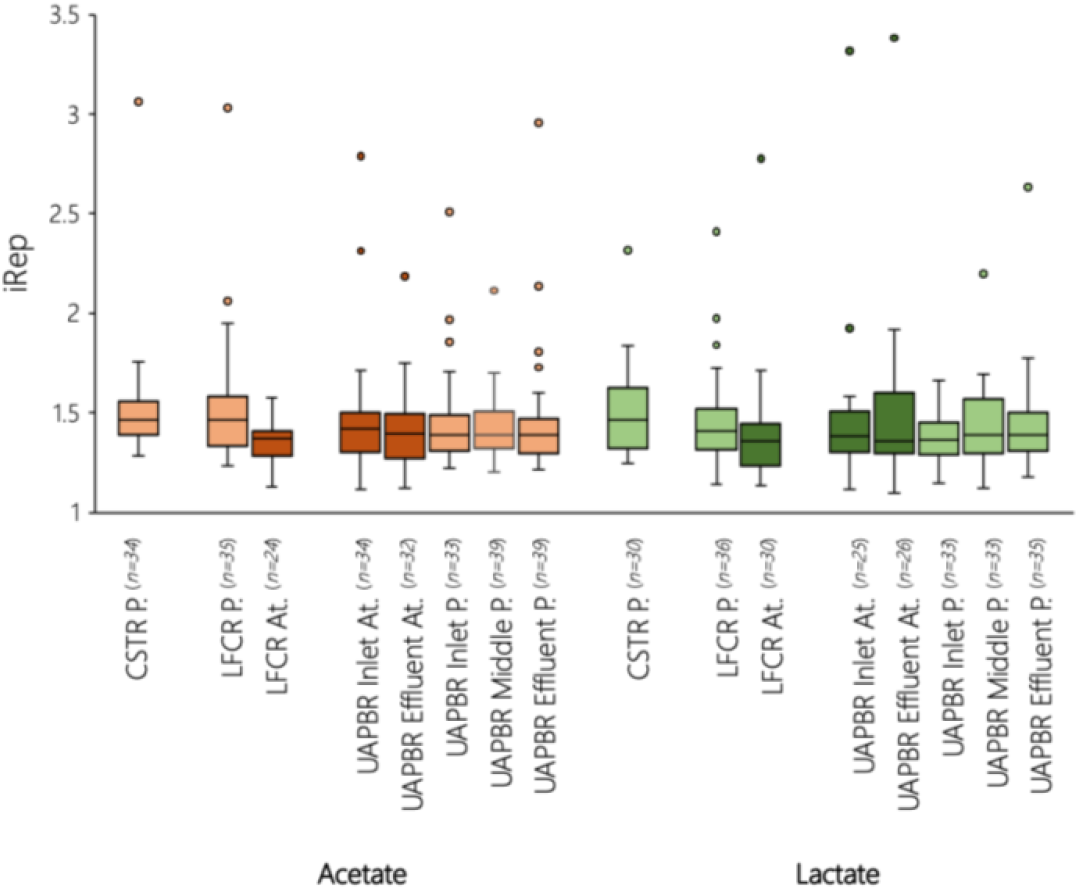
Indices of replications were used to characterise microbial replication rates of microorganisms in planktonic (P.) and attached biofilm (At.) communities. The interquartile range of iRep values determined for all microorganisms present within a reactor community is shown as box and whisker plots (*n* = number of iRep determined).

### Metabolic analysis of recovered genomes

Twenty-two of the recovered genomes were for organisms predicted to be capable of dissimilatory sulfate reduction (Figure 1), based on the recovery of genes encoding adenylylsulfate reductases, sulfate adenylyltransferases, and dissimilatory sulfite reductases. Twenty-one of these SRB were classified as Deltaproteobacteria, and a single genome classified to the Clostridial genus *Desulfomaculatum*. Each of the 22 genomes annotated as SRB encoded a group 1 NiFe hydrogenase, indicating their capacity to perform H_2_ oxidation, and many encoded additional NiFe and FeFe hydrogenases. The capacity to perform carbon fixation was found in six SRB genomes, five of which encoded the Wood-Ljungdahl pathway. The sixth, the genome of a *Desulfovibrio desulfuricans*, encoded the recently demonstrated autotrophic reductive glycine pathway which uses a subset of the Wood-Ljungdahl pathway and several enzymes involved in glycine reduction (Song et al., 2020; Sánchez-Andrea et al., 2020). In the genomes not linked to organisms carrying out dissimilatory sulfate reduction, hydrogen-evolving NiFe group 4 or FeFe hydrogenases were highly prevalent. A wide variety and broad distribution of hydrogenases has been described previously as a strong indication that hydrogen metabolism is important in an environment (Greening et al., 2016; Probst et al., 2017).

One further microorganism’s genome encoded the Wood-Ljungdahl pathway. This Clostridiales was the only microorganism which could be confirmed as a syntrophic acetate oxidising bacteria (SAOB) based on encoded gene pathways. However, 18 genomes of non-SRB encoded formate dehydrogenases, methylenetetrahydrofolate dehydrogenase (EC:1.5.1.5) and formate tetrahydrofolate ligase (EC:6.3.4.3) of the Wood-Ljungdahl pathway (but lacked the remainder of this pathway) together with a glycine cleavage pathway. This combined pathway was suggested as a possible mechanism of syntrophic acetate oxidation by Nobu et al. (2015). Thirteen of these potential SAOB also encoded enzymes necessary for citrate oxidation. These combined pathways may represent a possible mechanism for the oxidation of citrate to CO2 and H_2_. Hagen et al., (2016) proposed a similar mechanism of complete oxidation of fatty acids by a Firmicute, found in an anaerobic bioreactor, based on metagenomic and proteomic data.

The metabolic potential of the 50 most abundant genomes across the 17 metagenomes is presented in Figure 3. Nitrogen metabolisms were not predominant in the recovered genomes. The large majority of these genomes encoded glutamine synthetase (EC 6.3.1.2) or a glutamate dehydrogenase (EC:1.4.1.2) for the assimilation of ammonia, which was supplied in the reactor medium at 18.7 mM. In contrast, few genomes encode genes for denitrification, dissimilatory nitrate reduction and nitrification. However, nitrogenases (*nifDHK*) involved in nitrogen fixation were present in many of the SRB genomes and those of some Proteobacteria and Bacteroidetes. Fermentation pathways and genes needed for oxidation of various volatile fatty acids and several amino acid groups were common in the recovered genomes.

**Figure 3.**
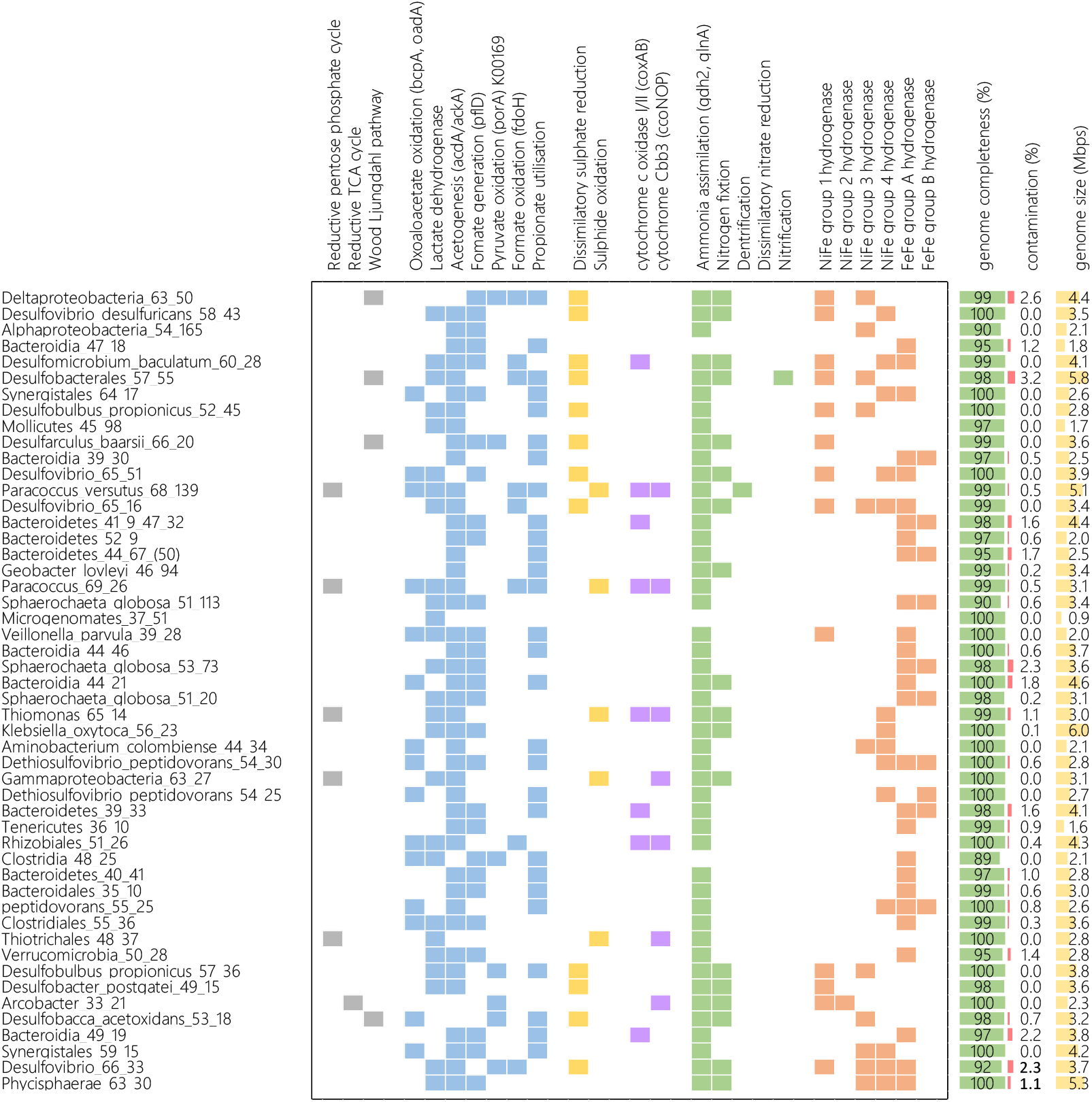
Genome statistics and metabolic features encoded by the genomes of the most prominent microorganisms associated with the BSR reactor systems and inoculum. Genome completeness and contamination were determined using CheckM. The shown metabolic features are described in detail in the supplementary data.

The majority of the recovered genomes could be divided into three broad groups based on encoded metabolisms (Figure S5). These genomes were constituted of SRB, of mainly fermentative microorganisms encoding FeFe group A hydrogenases, and a third group largely made up of genomes encoding oxidative phosphorylation electron transport chains including cytochrome c oxidases (caa3-type). The absence of gene pathways in these metagenomes relating to other forms of chemolithotrophy further indicates that fermentation, and respiration using sulphate and potentially trace oxygen are the predominant metabolisms in these reactor systems.

### Recovered hydrogenases

A total of 321 hydrogenases were identified in 130 genomes (Figure 4; Table S2), with 77 of these hydrogenases encoded by the 22 SRB. Each of these SRB typically encoded either a group 1a or 1b NiFe hydrogenase, groups of hydrogenases which are responsible for H_2_ uptake and are well-studied in SRB (Fauque et al., 1988). Group 3b and 3d NiFe hydrogenases were also common in the SRB genomes. These hydrogenases are responsible for electron bifurcation coupled to NADPH and bidirectional hydrogen activity coupled to NADH, respectively. Pyruvate ferredoxin oxidoreductases (*porA*), which catalyse the oxidation of pyruvate linked to the reduction of ferredoxin needed for hydrogen evolution, were also common in SRB. Many *Desulfovibrio* genomes and the recovered *Desulfomicrobium baculum* genome encoded group A FeFe hydrogenases and many of these SRB encoded NiFe group 4e hydrogenases, bidirectional hydrogenases which couple ferredoxin oxidation to H^+^ reduction (Greening et al., 2016). This is indicative of these microorganism’s capacity to contribute to H_2_ evolution. This dual metabolic capacity allows SRB to transition from sulfidogenic to hydrogenotrophic metabolism in the absence of sulfate (Plugge et al., 2011).

**Figure 4.**
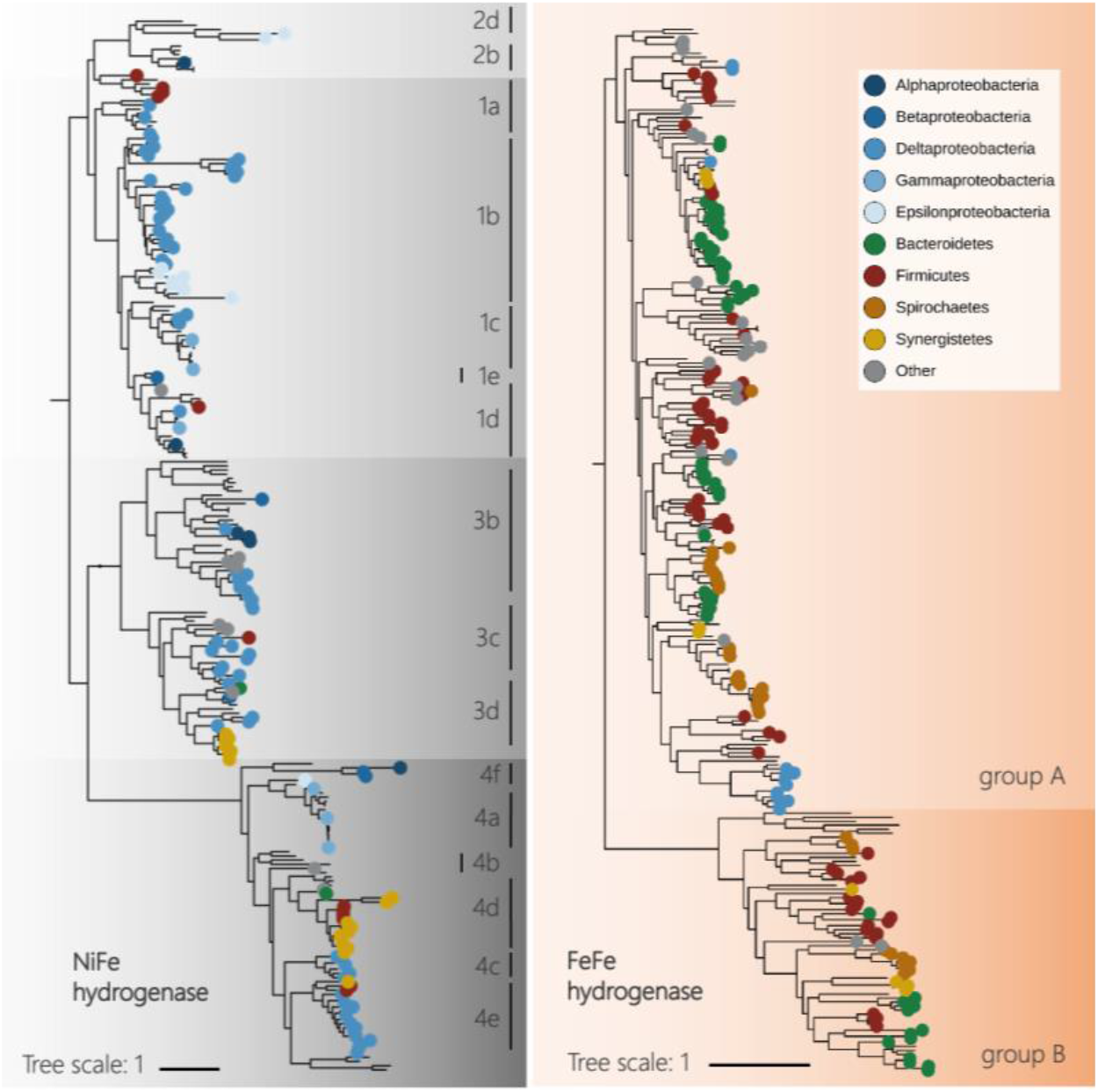
Phylogeny and classification of the 321 NiFe and FeFe hydrogenase protein sequences identified from 130 of the total 163 genomes recovered from the bioreactors of this study. Reference sequences are shown without phylum annotation. Hydrogenase genes were identified using custom-built HMMs and were further classified according to HydDB (Søndergaard et al., 2016). FeFe group A subgroups cannot be resolved phylogenetically but were further classified based on the immediately downstream catalytic subunit. Detailed classification of all recovered hydrogenases can be found in the supplementary data (Table S2).

Group 1 NiFe hydrogenases were identified in 26 Proteobacteria, 5 Firmicutes and a single Chlorobi genome. Hydrogen-evolving group 4 NiFe hydrogenases were found in Synergistales genomes (Figure 4). For example, group 4d NiFe hydrogenases, in which hydrogen evolution is linked to ferredoxin oxidation and sodium translocation (McTernan et al., 2014), were identified in 12 of the 15 recovered Synergistales genomes. These genomes commonly encoded additional FeFe hydrogenases and the bidirectional NADH-linked NiFe group 3d hydrogenases (Figure 4). Each of these genomes encoded a number of genes relating to amino acid catabolism, acetogenesis and oxaloacetate decarboxylases necessary for anaerobic citrate consumption (Bott, 1997), whilst none encoded any lactate dehydrogenases. One of these microorganisms, Synergistales_64_17, was dominant across most of the bioreactor communities, particularly the acetate- and the lactate-supplemented CSTRs. Pyruvate ferredoxin oxidoreductases (*porA*) implicated in hydrogen production were also common in Synergistetes genomes. Thus, we conclude that Synergistales bacteria are important in hydrogen production in the bioreactors, likely coupled to fermentation of amino acids and possibly citrate.

The capacity to produce hydrogen was predicted from the genomes of many other bacteria. Within 88 of the recovered genomes, almost all of which were classified as Bacteroidetes, Spirochaetes or Firmicutes (Figure 4), 182 FeFe hydrogenases were identified. FeFe hydrogenases are most commonly associated with hydrogen evolution (Vignais and Colbeau, 2004; Peters et al., 2015). The most commonly encoded were the trimeric group FeFe group A3 hydrogenase which reversibly bifurcates electrons between H_2_, and ferredoxin and NAD (Calusinska et al., 2010). Genes encoding Rnf complexes were common in these genomes. Pyruvate ferredoxin oxidoreductases (*porA*) implicated in hydrogen production were also present in some Firmicute genomes. Hydrogen can be generated by formate oxidation catalysed by a NiFe hydrogenase (Sawers, 1994) or group A4 FeFe hydrogenases (Calusinska et al., 2010) coupled to a formate dehydrogenase (Fdh), however, formate dehydrogenases were typically found in low-abundance genomes.

Overall, we found abundant evidence for H_2_ production and consumption in a diversity of bacteria across the reactors studied, including both those supplemented with lactate and acetate plus a small amount of fermentable substrate. Synergistales were abundant regardless of the substrate supplied, probably because they fermented amino acids and citrate common to all reactors. In general, the lactate reactors had more abundant and less diverse SRB. The absence of genes for chemolithotrophy other than in dissimilatory sulphate reducing bacteria, together with the physiochemical evidence of lactate, citrate and amino acid oxidation, suggests that nearly all the H_2_ which was generated in the reactors resulted from fermentation.

### Predicted metabolism of bacteria not involved in H_2_ metabolism

In genomes encoding no identified hydrogenases, partial and complete oxidative phosphorylation electron transport chains were common. These genomes, largely classified to Proteobacteria, encoded cbb3 oxygen reductases (Preisig et al., 1996) and bd-quinol oxidases (D’Mello et al., 1996) which have both been characterised with a high affinity for oxygen. However, functionally, the heme-copper cbb3 oxygen reductase would be inhibited by the hydrogen sulfide (Wu et al., 2015), which is present in the reactors at up to 330 mg/L. Thus, aerobic metabolism reliant on these reductases is unlikely in these reactors. However, some of these Proteobacteria may have consumed trace oxygen present in the medium using encoded sulfide-resistant bd-quinol oxidases (Forte et al., 2016). This functionality is important because influent solutions in real world applications contain trace O2 present, and its removal is beneficial to reactor performance. Some of these Proteobacteria also encoded sulfide (*sox*) gene pathways indicating trace oxygen consumption could be linked to the oxidation of the sulfide produced by SRB.

### UAPBRs’ microbial communities

Stratification of the microbial communities between the successive zones of the lactate-supplemented UAPBR occurred due to the plug-flow fluid dynamics that govern this reactor configuration (Figure S3). The performance of this system is summarised in Figure 5B and has been previously reported by Hessler et al. (2018). Briefly, the sulfate consumed within this reactor decreased between each subsequent reactor zone, from approximately 6.0 mM in the inlet zone to 0.6 mM in the effluent zone. This brought the sulfate conversion achieved by the overall reactor to 96%, at a four-day HRT. The sulfate reduced in each zone was putatively linked to the oxidation of various VFAs as shown in Figure 5B.

**Figure 5.**
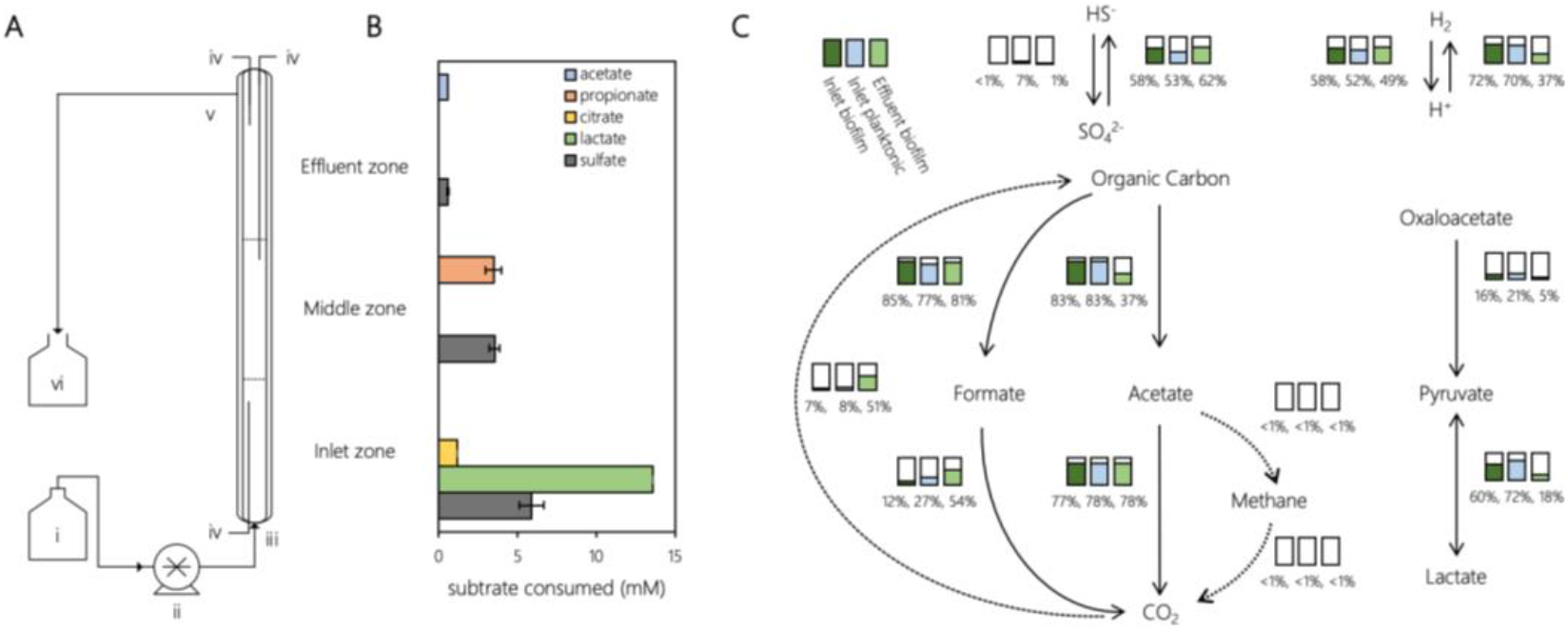
Schematic diagram (A) of the UAPBR showing the (i) feed reservoir, (ii) peristaltic pump, (iii) reactor inlet port, (iv) sampling ports, (v) effluent port, (vi) effluent collection, and the inlet, middle and effluent zones The amount of sulfate and of various VFAs which were confirmed to have been consumed (B) in the sequential zones of the lactate-supplemented UAPBR at a four-day HRT steady state. These are adapted from Hessler et al. (2018). (C) Metabolic reactions predicted from the genomes recovered from the biofilm and planktonic microbial communities of the lactate supplemented UAPBRs. Bar graphs and corresponding percentages represent the relative abundance of microorganisms within a community capable of performing a given reaction. The representation of microorganisms capable of performing autotrophy is elevated in the effluent attached community. Formate oxidation and carbon fixation through the Wood-Ljungdahl pathway are difficult to separate as formate dehydrogenase is a key enzyme in this autotrophic pathway. Of the 51% of the effluent biofilm community capable of carbon fixation, 41% encoded the Wood-Ljungdahl pathway, which is therefore represented in the proportion of microorganisms encoding a formate dehydrogenase.

Genome-resolved metagenomic analysis of the microbial communities in the lactate-supplemented UAPBR reactor revealed substantial differences in the composition of the biofilm microbial communities and their metabolic features between the inlet and effluent zones (Figure S3). Changes in the planktonic phase were less pronounced, likely a result of the plug flow carrying the planktonic community throughout the three zones irrespective of individual microbial activity. The inlet zone biofilm communities were dominated by SRB and fermentative microorganisms capable of lactate oxidation. The proportion of the microbial communities of the inlet zone’s biofilm and planktonic community which encoded a lactate dehydrogenase decreased from 60 and 72%, respectively, to 18% of the effluent biofilm community (Figure 5C). This corresponded with the representation of genomes encoding genes for acetogenesis decreasing from over 80% in both the inlet zone biofilm and planktonic community to 37% in the biofilm of the effluent zone. The representation of oxaloacetate decarboxylases (EC:7.2.4.2 and EC:4.1.1.112), enzymes involved in anaerobic citrate consumption (Bott, 1997), were far greater in the inlet zone communities than the biofilm community of the effluent zone. These features correlate with the restriction of citrate and lactate oxidation to the inlet zone.

The representation of group 1 NiFe hydrogenases was consistent across the lactate-supplemented UAPBR communities due to consistent representation of SRB across these communities. However, the representation of hydrogen evolving hydrogenases decreased substantially between the inlet and effluent zone. Carbon fixation gene pathways and formate dehydrogenase genes were enriched in the biofilm community of the effluent zone. However, these carbon fixation pathways were largely constituted by three SRB genomes, namely Deltaproteobacteria_63_50, Desulfarculus_baarsii_66_20 and Deltaproteobacteria_58_15, that each encoded the Wood-Ljungdahl pathway. This accounts for the increased proportion of formate dehydrogenases, a key enzyme in this autotrophic pathway, in this community. The Wood-Ljungdahl pathway may likely have been operated in the oxidative direction in these SRB for the consumption of acetate, which was observed in this zone of the reactor.

The predicted autotrophic Deltaproteobacteria_63_50 was also prevalent within the biofilm in the effluent zone of the acetate-supplemented UAPBR. Its dominance in the effluent biofilm communities of both acetate and lactate UAPBRs, but low abundance in these reactors inlet zones, likely indicates this SRB’s capacity to compete for sulfate at low concentrations (<100 mg/L). The effective cultivation of this organism within a bioreactor community may confer enhanced sulphate conversions under similar conditions.

The high sulphate conversions achieved by the lactate amended UAPBR was previously attributed to the out-competition of fermentative microorganisms for lactate by SRB in the inlet zone, and secondly the zoning of the reactor allowing different SRB suited to the oxidation of different electron donors to effectively compete in niche environments along the length of the reactor (Hessler et al., 2018). What was not considered in previous analyses was the effect of hydrogen produced during fermentation of lactate, yeast extract and citrate in the inlet zone, on the growth of SRB present in the middle and effluent zones. It is apparent from the wide distribution and the diversity of recovered hydrogenases in this reactor that hydrogen cycling is an important feature in these microbial communities. This suggests that the maintained sulfate reduction rates observed in the effluent zone, at low sulfate concentrations, were enabled by consumption of hydrogen in addition to acetate by SRB in this zone.

Kinetic modelling of the sulfate reduction reactions observed in the primarily acetate-supplemented UAPBR over the course of an HRT study found the rate of sulfate reduction in the reactor could be described with a uniform reaction order of 2.9 (unpublished). This high reaction order was not anticipated, as acetate-oxidising SRB are typically characterised with very high affinities for both acetate and sulfate (Ingvorsen et al., 1984; O’Flaherty et al., 1998). Genome-resolved metagenomic analyses indicate that H_2_ is likely to have been produced through fermentation and subsequently consumed by SRB in these reactors and, therefore, may explain this elevated reaction order.

### Acetate-supplemented CSTR

The acetate-CSTR maintained sulfate-reduction at far greater dilution rates than had been anticipated based on a previous study using an acetate-supplemented CSTR and similar operating conditions. This study, conducted by Moosa et al. (2002), however, did not include yeast extract nor citrate in the media. This bioreactor without fermentable substrates exhibited washout of SRB at a dilution rate of 0.021 h^-1^ (two-day HRT), whereas the CSTR from our study was able to sustain sulfate-reduction at a 0.042 h^-1^ dilution rate (one-day HRT; Figure 6A) and was able to achieve greater VSRR.

**Figure 6.**
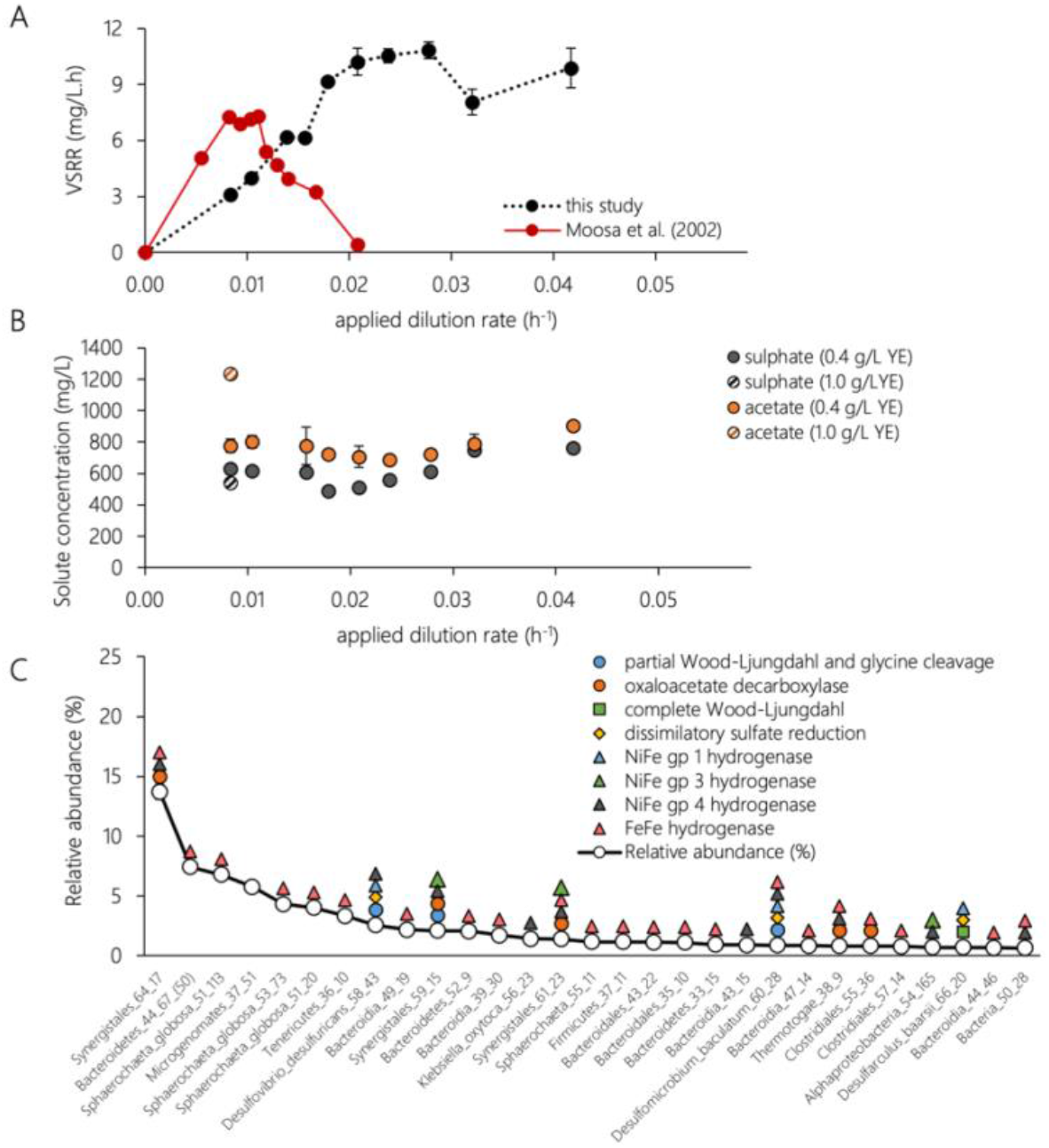
A) The performance of this CSTR is shown in terms of the achieved volumetric sulfate reduction rate (VSRR) at increasing volumetric sulfate loading rates (VSLR), and shown against a similarly operated CSTR (Moosa et al., 2000) which was not supplemented with yeast extract nor citrate. B) Steady-state residual acetate and sulfate concentrations in the acetate-supplemented CSTR at a range of applied dilution rates. The concentration of yeast extract in the medium was decreased from 1.0 g/L to 0.4 g/L before the HRT study was begun which allowed the amount of acetate generated through yeast extract oxidation to be estimated. The residual acetate concentration in the reactor was then predicted based on the assumption that all sulfate reduced was linked to acetate oxidation and that the 0.4 g/L yeast extract further contributed to the residual acetate concentration.. Error bars of performance data represent one standard deviation from the mean (*n*>4). C) The CSTR microbial community at a four-day HRT is shown as an annotated rank abundance curve showing various genome inferred metabolic features. Error bars represent one standard deviation from the mean of two independently sampled metagenomes.

At the five-day HRT steady-state, the yeast extract concentration in the media was reduced from 1.0 g/L to 0.4 g/L. This led to a decrease in the residual acetate concentration in the reactor and allowed the total amount of acetate produced through the oxidation of the remaining 0.4 g/L yeast extract to be estimated to 268 mg/L (Figure 6B). Therefore, the acetate available in the feed (720 mg/L) and produced from yeast extract oxidation (268 mg/L) could be estimated at 988 mg/L. The reduction in the yeast extract concentration by 600 mg/L led to a minimal increase in residual sulfate from 537 mg/L to just 630 mg/L, indicating that yeast extract was not the predominant electron donor linked to sulfate reduction. Instead, we attribute the increased sulfate concentration to the decrease in hydrogen produced through fermentation. The residual acetate concentration in the reactor varied between 688 and 904 mg/L during the HRT study. The stoichiometric oxidation of acetate by SRB largely accounts for the degree of acetate oxidation at each HRT. However, citrate nor any predictable by-products of its oxidation were detected in the reactor at any steady-state. Whether this citrate was sequentially fermented and the organic products oxidised by separate microorganisms to CO2 or whether citrate was oxidised completely to CO2 within individual microorganisms remains unclear.

Metagenomic analysis of the CSTR community at the four-day HRT found that the most abundant bacteria in the reactor had fermentation-based metabolisms (Figure 6C). Several Synergistales bacteria are predicted to be capable of oxidation of the citrate, with one of these Synergistales also encoding the partial Wood-Ljungdahl and glycine cleavage pathway predicted to allow the oxidation of acetyl-CoA to CO2 and H_2_. The community was dominated by a Synergistales (Synergistales_64_17), as well as one Bacteroidetes and three Spirochaetes. All of the genomes of these abundant bacteria encoded hydrogen-evolving FeFe hydrogenases. Each of these five genomes also contained a variety of genes relating to amino acid metabolism (Figure S4). Aspartate/tyrosine/aromatic aminotransferase (EC:2.6.1.1, EC:2.6.1.57), Branched-chain amino acid aminotransferase/4-amino-4-deoxychorismate lyase (EC:2.6.1.42, EC:4.1.3.38) and Histidinol-phosphate/aromatic aminotransferase (EC:2.6.1.9) were represented in approximately 70% of the genomically defined microbial community, of which 56% were from Bacteroidetes, Spirochaetes or Synergistales. The dominance of these fermentative microorganisms is therefore thought to be afforded through the utilisation of yeast extract and possibly citrate, with hydrogen likely being used as the terminal electron acceptor.

Three predominant SRB were identified in this CSTR community (Figure 6C). These were classified as *Desulfomicrobium baculatum*, *Desulfovibrio desulfuricans* and *Desulfarculus baarsi*. These three SRB were widespread in each of the six reactors, with *D. baculatum* and *D.desulfuricans* particularly prevalent in both acetate- and lactate-supplemented planktonic communities. Each of these genomes encoded either a full or partial Wood-Ljungdahl pathway with a glycine cleavage pathway, as well as acetyl-CoA synthetases (EC:6.2.1.1), indicating their likely capacity for acetate oxidation. No methanogens were detected and only one potential SAOB was found, a Synergistales encoding a partial Wood-Ljungdahl and glycine cleavage pathway. This supports the conclusion that consumed acetate was the primary electron donor for sulfate reduction. Each of these SRB genomes also encoded group 1 NiFe hydrogenases indicating their capacity for H_2_ utilisation. Fermentative hydrogen-generating microorganisms rely on other anaerobes to maintain low hydrogen partial pressures to allow fermentation to remain energetically favourable (Stams et al., 2006; McInerney et al., 2008). The absence of methanogens and the positioning of the SRB in this community, with nearly ubiquitous distribution of hydrogen-evolving hydrogenases amongst the non-SRB suggests these SRB are likely to be consuming hydrogen generated by these fermentative microorganisms whilst sourcing carbon predominantly from acetate. This simultaneous hydrogen and acetate consumption by SRB has been documented in pure and co-culture experiments (Sorokin, 1966; Jansen et al., 1984) and the supplementation of acetate-supplemented sulfidogenic cultures with hydrogen has been suggested by Stams and Plugge (2009) to allow better enrichments of SRB. These results indicate that the VSRR of acetate-supplemented BSR reactors could be enhanced through supplementation of low concentrations of fermentable material to support a co-existing fermentative, hydrogen-evolving, microbial consortium.

## Conclusion

BSR represents a promising strategy for the low-cost bioremediation of ARD yet a consensus on the operating conditions required for robust reactor operation and efficient utilisation of the supplied electron donor(s) for such a process has not yet been reached. These discussions are seldom informed by the metabolic potential or the microbial composition held within these bioreactors. Here, we described the genome-resolved metagenomic characterisation of the microbial communities associated with six well-functioning continuous BSR reactor systems. This approach allowed the reconstruction of 163 genomes of microorganisms associated with this process. Within these metagenomes we identified 321 hydrogenases, classified into several groups of NiFe and FeFe hydrogenases. Evaluation of these hydrogenases and co-occurring metabolic features indicated that hydrogen was an important energy carrier in these communities. We were able to postulate that the acetate-oxidising SRB in these reactor systems benefitted from the hydrogen generated by a range of microorganisms encoding fermentative metabolisms. The fermentative microorganisms in these reactors were sustained through fermentation of yeast extract, lactate and possibly citrate. We conclude that recreating and further manipulating the conditions observed in these reactors to stimulate hydrogen evolution through fermentation should lead to the enhanced growth of acetate-oxidising SRB and consequently enhanced reaction rates and successful operation at higher dilution rates. This approach would allow acetate, a low-cost electron donor, to become more viable for ARD remediating BSR processes.

## Methodology

### Microbial culture

The six reactor systems were inoculated with a mixed sulfidogenic microbial culture described by Hessler et al (2018). This microbial culture was maintained, prior to inoculation, on neutral modified Postgate B medium (0.42 g/L KH_2_PO_4_, 1.0 g/L NH_4_Cl, 1.0 g/L MgSO_4_.7H_2_O, 0.9 g/L Na_2_SO_4_, 0.4 g/L yeast extract, 0.3 g/L sodium citrate) supplemented with various electron donors including acetate, lactate, ethanol and algal anaerobic digestate.

### Reactor configurations and operation

Three lab-scale bioreactor configurations were operated in this study, namely continuous stirred tank reactors (CSTR), linear flow channel reactors (LFCRs) and up flow anaerobic packed bed reactors (UAPBRs). Each of these reactor systems was operated in duplicate and continuously fed the modified postgate B medium supplemented with either sodium acetate (0.92 g/L) or sodium lactate (1.2 g/L) as the electron donor. The CSTR and UAPBR configurations are described by Hessler et al. (2018) and the LFCR is described by Hessler et al. (2020a).

### Analytical methods

The sulfide produced and the residual sulfate in the reactors were quantified using the DMPD (Cline, 1969) and APHA (1975) turbimetric methods (Greenberg and Eaton, 1999), respectively. The residual and generated volatile fatty acids in the reactor, including acetate, lactate, propionate, citrate, butyrate and valerate, were quantified as described by Hessler et al. (2018). The pH was monitored using a Cyberscan 2500 micro pH meter fitted with an XS Sensor 2-Pore T DHS pH probe and the redox potential of each sample was measured using the Metrohm 827 pH lab meter fitted with a Pt-ring KCl electrode (Metrohm model 6.0451.100).

### Steady-state biological sampling

Biofilm and planktonic cells were isolated for metagenomic sequencing and cell quantification from the reactors at steady-state at a four-day HRT. Solid support structures were removed from the LFCR and UAPBR using sterile stainless-steel scissors and forceps. The biofilm associated cells were recovered through gentle agitation in reactor medium. Cells embedded in the biofilms attached to the solid support structures were isolated in duplicate for cell quantification using a non-destructive whole cell detachment protocol described by Hessler et al. (2018). The cells from each of these defined phases were quantified, in duplicate, through direct cell counting oil immersion phase contrast microscopy (Olympus BX40) and a Thoma counting chamber. Recovered planktonic and biofilm-associated cells were recovered for total DNA extraction by centrifugation at 10 000 g for 10 min. Total DNA extraction of biofilm attached cells was performed directly from the colonised solid support structures.

### Metagenomic DNA extraction and sequencing

Total genomic DNA was extracted from 34 metagenomic samples, representing the 16 bioreactor and the inoculum sample, in duplicate, using a NucleoSpin^®^ Soil Genomic DNA extraction kit (Machery-Nagel, Germany) as per manufacturer’s instructions. Library preparation and Illumina^®^ sequencing was performed at UC Berkeley’s Functional Genomics Laboratory. Paired-end Illumina^®^ sequence libraries were prepared with an insert size of 400-800 bp and were sequenced on an Illumina^®^ HiSeq4000^®^ generating 2.5 Gbp of raw data per sample.

### Read processing, assembly and annotation

All read sets were trimmed using Sickle (https://github.com/najoshi/sickle) using default parameters. Sequencing reads originating from metagenomes sampled in duplicate were concatenated into single files and were processed in parallel with the two original un-concatenated read sets. Reads were assembled using MEGAHIT version 1.1.3 (Li et al., 2015) with the following parameters: --k-min 21, --k-max 99, --k-step 10, --min—count 2. Read mappings for all scaffolds were determined using Bowtie2 (Langmead and Salzberg, 2012). Open reading frames (ORFs) were predicted using Prodigal’s metagenome procedure (Hyatt et al., 2010). These ORFs were then annotated using USEARCH (Edgar, 2010) against KEGG (Kanehisa and Goto, 2000), UniRef100 and Uniprot (Consortium, 2009) databases. Functional genes were predicted using Hmmsearch (http://hmmer.org/) against in house Hidden Markov Models (HMM) built from KEGG orthology groups, TIGRFAM and PFAM databases (found at https://github.com/banfieldlab).

### Hydrogenase classification and phylogeny

The protein sequences of the catalytic FeFe and NiFe hydrogenase subunits were identified using HMMs described above and were Muscle aligned v3.8.31 (Edgar, 2004) and trimmed in Geneious v2019.0.4. Maximum-likelihood phylogenetic trees for NiFe and FeFe hydrogenase protein sequences were constructed using Fasttree2 (Price et al., 2010). The hydrogenases were classified based on the best-hit HMM and the presence of consensus motifs surrounding the conserved cysteine residues and further verified using HydDB (Søndergaard et al., 2016). Protein sequences classified as group A FeFe hydrogenases were further classified to group A1, A2, A3 or A4 based on the downstream catalytic subunit protein sequence (Søndergaard et al., 2016).

### Binning of metagenomes

Individual binning was performed on each of the 17 concatenated and 34 unconcentatented assemblies using MaxBin (Wu et al., 2014), Metabat (Kang et al., 2015) and Concoct (Alneberg et al., 2014) and manual binning performed on the basis of GC content, coverage and consensus taxonomy using ggKbase (ggkbase.berkeley.edu). Series coverage information across the 51 samples was used for binning in parallel using Concoct. A set of non-redundant genome bins were generated for each metagenome using Das Tool (Sieber et al., 2018) and a non-redundant set of genome bins across the 51 assemblies was selected using dRep (Olm et al., 2017). The degree of genome completeness and contamination was determined using CheckM (Parks et al., 2015) based on the recovery of bacterial and archaeal single copy genes. This was repeated using checkm’s CPR specific gene marker set for the evaluation of a recovered Microgenomates genome bin.

### Indices of replication

Relative instantaneous microbial growth rates were estimated using Indices of replication (iRep). These were calculated using iRep.py (Brown et al., 2016), using default settings, for all genomes which passed iRep’s quality threshold.

### Phylogenetic analyses

Phylogenetic analyses were performed using 16 universal ribosomal proteins. Sixteen ribosomal protein sequences from 2896 reference organisms, as described by Hug et al. (2016), and each of the recovered genome bins were aligned using MUSCLE v3.8.31 (Edgar, 2004), trimmed using Geneious v2019.0.4. and subsequently concatenated into a single alignment. A maximum-likelihood Phylogenetic reconstruction of these two alignments were performed using FastTree2 (Price et al., 2010).

### Hierarchical clustering and principal component analysis

Hierarchical clustered heatmaps and PCA analyses were produced using ClustVis (Metsalu and Vilo, 2015). Hierarchical clustering was performed using Ward’s method and Euclidean distances applied to ln(x+1) transformed values. Nonlinear iterative partial least squares (Nipals) PCA was used to assess the encoded metabolic genes across genome bins.

## Supporting information

Supplementary materials (KEGG summary)

## Acknowledgements

This study was funded through the Water Research Commission (K5-2393) and the DST/NRF of South Africa through Prof. Harrison’s SARChI Chair in Bioprocess Engineering (UID 64778). Dr Huddy was funded through DST/NRF Competitive Support for Unrated Researchers (CSUR) Grant (UID 111713). Dr Hessler was funded through NRF Scarce-skills MSc scholarship (UID 108052). We thank Dr Shufei Lei, Katherine Lane, Dr Rohan Sachdeva, Dr Patrick West, and the QB3 Vincent J. Coates Genomics Sequencing Laboratory for research support.

## Data availability

The genomes and raw sequencing reads will be made available under the NCBI project number PRJNA728813. Genomes will also be made available at https://ggkbase.berkeley.edu/Biological_sulphate_reduction_analysis/organisms.

