## Supplementary materials (Tables and Figures) for "Syntrophic H_2_ production enhances the performance of primarily acetate-supplemented reactors treating sulphate contaminated solutions"

### Evidence of syntrophic hydrogen metabolism linked to enhanced performance in a range of sulfate reducing reactor systems.

T. Hessler[1], S.T.L. Harrison[1,2], J.Banfield[3,4,5,6], R.J. Huddy[1,2*]

[1]The Center for Bioprocess Engineering Research, University of Cape Town, South Africa
[2]The Future Water Institute, University of Cape Town, South Africa
[3]The Innovative Genomics Institute at the University of California, Berkeley, California, USA
[4]The Department of Earth and Planetary Science, University of California, Berkeley, California, USA
[5]The Department of Environmental Science, Policy and Management, University of California, Berkeley, California, USA
[6]The University of Melbourne, Victoria, Australia

#### Supplementary information

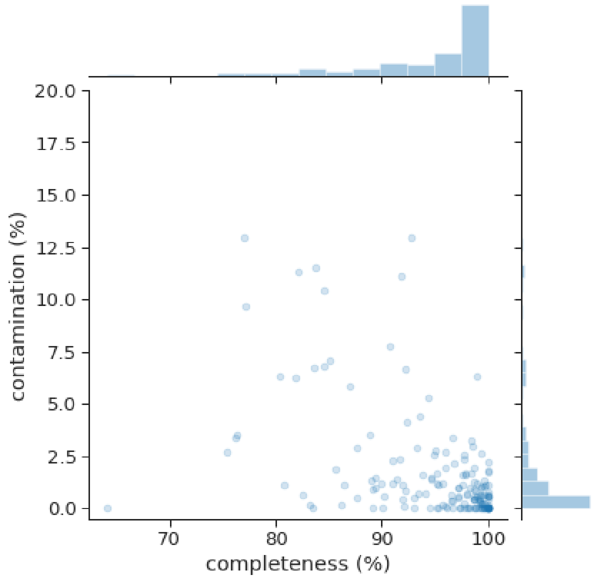


**Figure S1.** The quality of the MAGs reconstructed in this study shown as percentage contamination and completeness as determined by checkM based on the recovery of single copy genes.

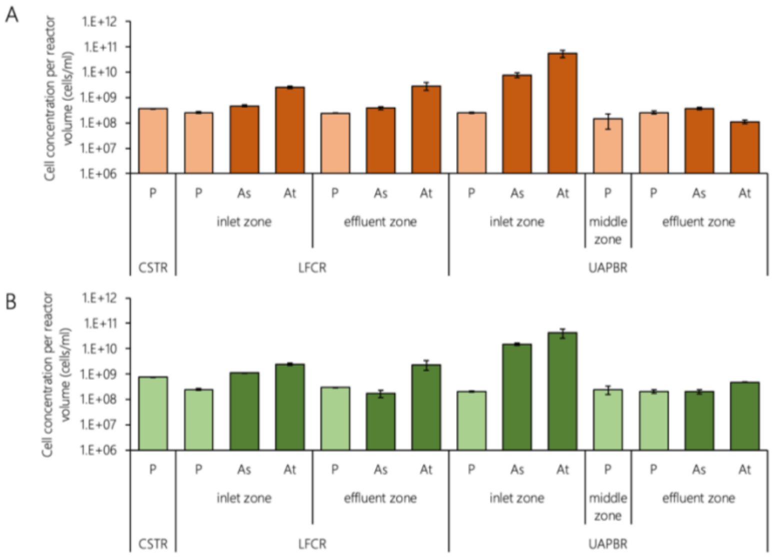


**Figure S2**. The concentration of planktonic, biofilm-associated and biofilm-attached phases retained within the (A) acetate and (B) lactate supplemented CSTRs, LFCRs and UAPBRs at steady state at a four-day HRT. Biofilm communities were isolated using a modified detachment protocol and all three phases were quantified through direct cell counting. Cell concentrations are normalised per total volume of reactor or reactor zone. Error bars represent one standard deviation from the mean. Samples were isolated in duplicate and counted in duplicate.

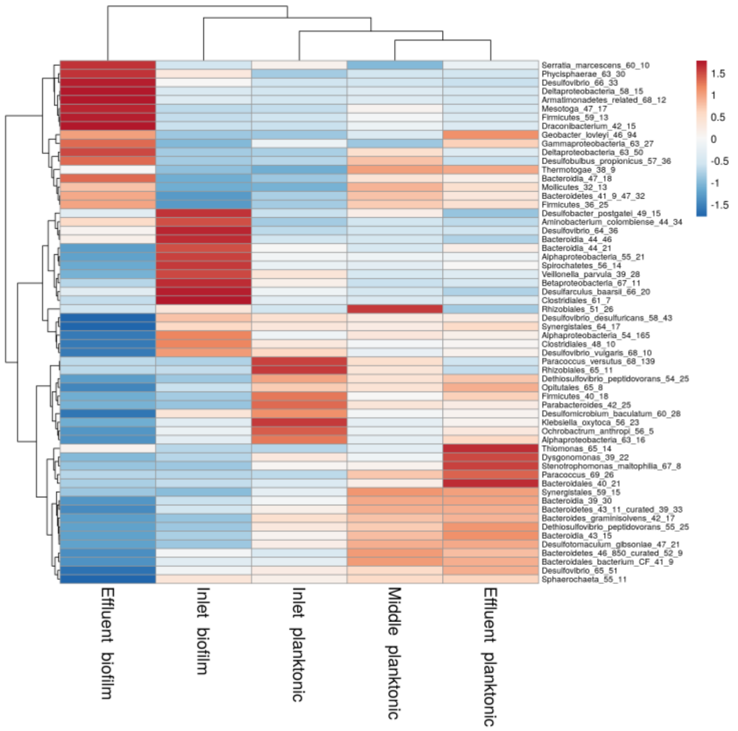


**Figure S3.** Hierarchical clustered heatmap showing the Z-scores of the 60 most abundant organisms across the biofilm and planktonic communities of the lactate-supplemented UAPBR inlet, middle and effluent zones.

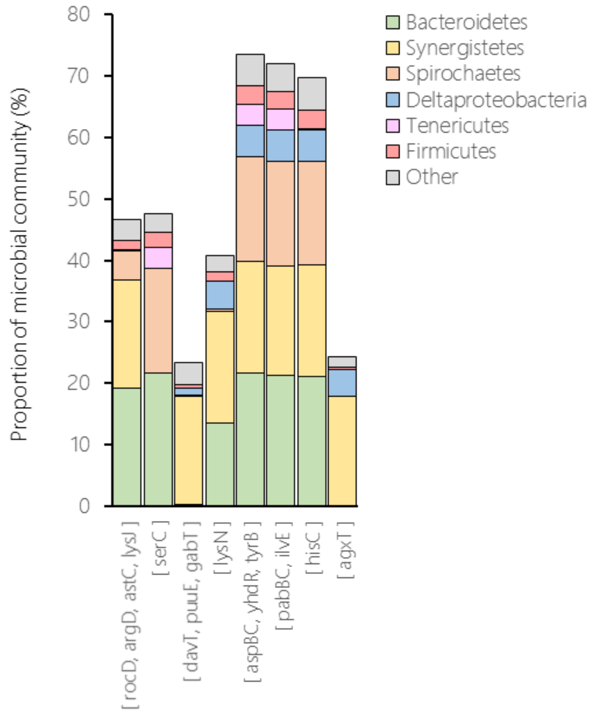


**Figure S4**. Proportion of the CSTR microbial community, and the taxonomic groups, encoding the following genes involved in amino acid utilisation: Ornithine/acetylornithine aminotransferase (*rocD*, *argD*, *astC*, *lysJ*), Phosphoserine aminotransferase (*serC*), 4-aminobutyrate aminotransferase and related aminotransferases (*davT*, *puuE*, *gabT*), Aminotransferase class I and II (*lysN*), Aspartate/tyrosine/aromatic aminotransferase (*aspBC*, *yhdR*, *tyrB*), Branched-chain amino acid aminotransferase/4-amino-4-deoxychorismate lyase (*pabBC*, *ilvE*), Histidinol-phosphate/aromatic aminotransferase (*hisC*) and Serine-pyruvate aminotransferase/archaeal aspartate aminotransferase (*agxT*). The classification of the genomes encoding these genes are shown in the legend.

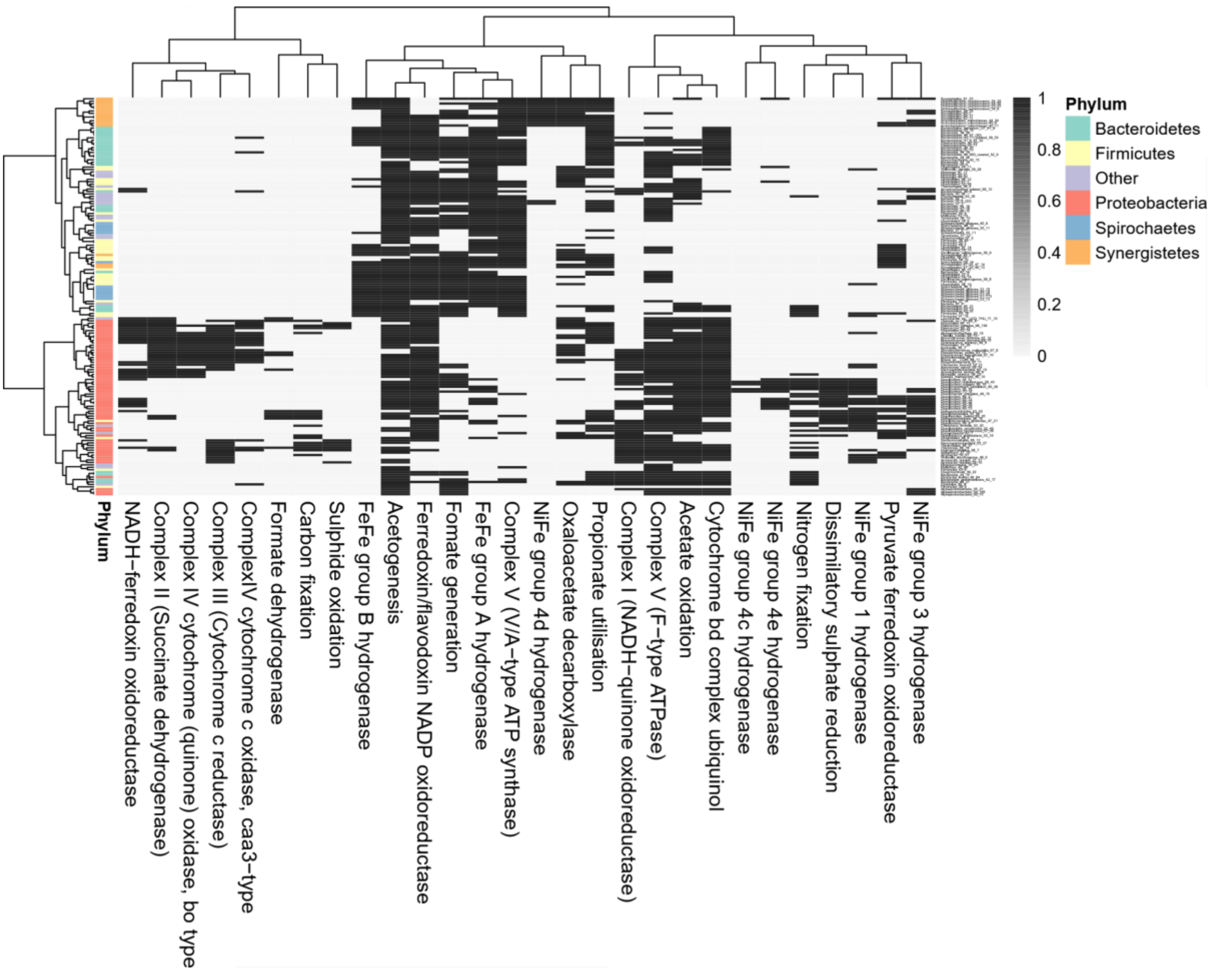


**Figure S5.** Hierarchical clustering of metabolic features encoded across the 163 recovered microbial genomes described in this study. The phylum to which each genome was classified is shown as row annotations. Clustering resolved these genomes, based on the selected genetic features, into two large clusters: the first representing genomes encoding largely fermentative metabolisms with FeFe hydrogenases, and a second representing genomes encoding alternative metabolisms such as oxidative phosphorylation, dissimilatory sulphate reduction, sulfide oxidation and carbon fixation with various NiFe group hydrogenases.

**Table S1.** Steady-state data collected from the six BSR reactors at a four-day HRT, at the time of total genomic DNA sampling. These data include pH and redox potential measurements, assayed sulfur and VFA concentrations and performance characteristics such as sulfate conversion and volumetric sulfate reduction rates (VSRR) at varying volumetric sulfate loading rates (VSLRs). The UAPBR is plug-flow governed reactor demarcated into three sequential zones of equal volume. The data collected from each of these zones is reported.

| | | Physiochemical measurements | | | | Solute concentrations (mg/L) | | | | | | | | | | | | | Reactor performance | | | | |
|---|---|---|---|---|---|---|---|---|---|---|---|---|---|---|---|---|---|---|---|---|---|---|---|
| reactor system | zone | ph | ± | redox potential (mV) | ± | sulfate | ± | sulfate reduced | sulfide | ± | citrate | ± | lactate | ± | acetate | ± | propionate | ± | VSLR (mg/L.h) | sulfate conversion | VSRR (mg/L.h) | ± | reference |
| acetate CSTR | n/a | 7.19 | 0.23 | -389.5 | 2.1 | 616.5 | 12.1 | 383.5 | 140.9 | 8.9 | 0.0 | 0.0 | n/a | n/a | 802.0 | 120.4 | n/a | n/a | 10.42 | 0.38 | 3.99 | 0.13 | |
| lactate CSTR | n/a | 7.06 | 0.03 | -384.5 | 2.1 | 430.6 | 10.1 | 569.4 | 195.9 | 10.0 | 0.0 | 0.0 | 0.0 | 0.0 | 745.7 | 120.4 | 161.5 | 12.0 | 10.42 | 0.57 | 5.93 | 0.11 | (Hessler et al., 2018a) |
| acetate LFCR | effluent zone | 7.51 | 0.00 | -379.0 | 18.4 | 312.4 | 27.7 | 687.6 | 212.2 | 27.9 | 0.0 | 0.0 | n/a | n/a | 643.0 | 50.4 | n/a | n/a | 10.42 | 0.69 | 7.16 | 0.29 | (Hessler et al., 2020) |
| lactate LFCR | effluent zone | 7.11 | 0.04 | -380.0 | 0.0 | 240.2 | 18.7 | 759.8 | 231.4 | 16.0 | 0.0 | 0.0 | 0.0 | 0.0 | 762.8 | 12.7 | 32.1 | 71.7 | 10.42 | 0.76 | 7.91 | 0.20 | (this study) |
| Acetate UAPBR | inlet zone | 7.38 | 0.07 | -341.7 | 2.1 | 217.1 | 34.1 | 782.9 | 274.6 | 11.4 | 0.0 | 0.0 | n/a | n/a | 558.3 | 74.0 | n/a | n/a | 31.25 | 0.78 | 24.47 | 1.07 | (Hessler et al., 2018b) |
| | middle zone | 7.40 | 0.09 | -344.3 | 1.5 | 54.4 | 21.7 | 162.7 | 367.7 | 36.9 | 0.0 | 0.0 | n/a | n/a | 477.1 | 31.9 | n/a | n/a | 6.78 | 0.75 | 5.08 | 0.68 | |
| | effluent zone | 7.41 | 0.08 | -351.0 | 1.0 | 33.8 | 19.0 | 20.6 | 339.5 | 45.6 | 0.0 | 0.0 | n/a | n/a | 477.1 | 31.9 | n/a | n/a | 1.70 | 0.38 | 0.64 | 0.59 | |
| | overall reactor | | | | | | | | | | | | | | | | | | 10.42 | 0.97 | 10.06 | 0.20 | |
| Lactate UAPBR | inlet zone | 7.09 | 0.01 | -332.0 | 8.5 | 433.1 | 74.4 | 566.9 | 204.7 | 23.5 | 0.0 | 0.0 | 0.0 | 0.0 | 719.4 | 28.6 | 191.0 | 36.9 | 31.25 | 0.57 | 17.71 | 2.32 | (Hessler et al., 2018a) |
| | middle zone | 7.15 | 0.08 | -343.0 | 5.7 | 93.3 | 31.7 | 339.9 | 319.4 | 18.6 | 0.0 | 0.0 | 0.0 | 0.0 | 781.5 | 37.1 | 0.0 | 0.0 | 13.54 | 0.78 | 10.62 | 0.99 | |
| | effluent zone | 7.22 | 0.05 | -348.0 | 4.2 | 37.8 | 6.8 | 55.4 | 304.1 | 46.7 | 0.0 | 0.0 | 0.0 | 0.0 | 764.8 | 79.3 | 3.8 | 9.6 | 2.91 | 0.59 | 1.73 | 0.21 | |
| | overall reactor | | | | | | | | | | | | | | | | | | 10.42 | 0.96 | 10.02 | 0.07 | |

**Table S2.** The classifications of the catalytic hydrogenase genes encoded by the microbial genomes recovered from the six BSR reactors and inoculum described in this study. The number of unique hydrogenase genes is shown per phylum. The function of the classified hydrogenase is shown in parentheses.

| | NiFe Group 1a (H2-uptake) | NiFe Group 1b (H2-uptake) | NiFe Group 1c (H2-uptake) | NiFe Group 1d (H2-uptake) | NiFe Group 1e (Bidirectional) | NiFe Group 1f (H2-uptake) | NiFe Group 2b (H2-sensing) | NiFe Group 2d (H2-uptake) | NiFe Group 3b (Bidirectional) | NiFe Group 3c (Electron-bifurcation) | NiFe Group 3d (Bidirectional) | NiFe Group 4a (H2-evolution) | NiFe Group 4b (H2-evolution) | NiFe Group 4c (H2-evolution) | NiFe Group 4d (H2-evolution) | NiFe Group 4e (Bidirectional) | NiFe Group 4f (unconfirmed) | FeFe group A unclassified | FeFe group A1 (H2-evolution) | FeFe group A2 (unconfirmed) | FeFe group A3 (Bidirectional) | FeFe group A4 (Bidirectional) | FeFe Group B (H2-evolution) |
|---|---|---|---|---|---|---|---|---|---|---|---|---|---|---|---|---|---|---|---|---|---|---|---|
| Synergistetes | | | | | | | | | | | 6 | | | | 12 | 1 | | | | | 8 | | 8 |
| Bacteroidetes | | | | | | | | | | | 1 | | | | | | | | 9 | | 33 | | 12 |
| Spirochaetes | | | | | | | | | | | | | | | | | | 1 | | | 21 | | 10 |
| Firmicutes | 4 | | | 1 | | | | | 1 | 1 | | | | | | 2 | | 3 | 3 | 7 | 25 | 1 | 15 |
| Atribacteria | | | | | | | | | | | | | | | 2 | | | 2 | | | | | |
| Microgenomates | | | | | | | | | | | | | | | | | | | | | | | |
| Lentishphaerae | | | | | | | | | | | | | | | | | | | | | 2 | | 1 |
| Tenericutes | | | | | | | | | | | | | | | | | | | | | 3 | | |
| Planctomycetes | | | | | | | | | 1 | | 1 | | 1 | | | | | | | | 5 | | |
| Chlorobi | | | | 1 | | | | | 1 | | | | | | | | | | | | | | |
| Actinobacteria | | | | | | | | | | | | | | | | | | | | | | | |
| Verrucomicrobia | | | | | | | | | | | | | | | | | | | | | 1 | | |
| Thermotogae | | | | | | | | | | | | | | | | | | 1 | 2 | | 3 | | 1 |
| Cloacimonetes | | | | | | | | | | | | | | | | | | | | | | | |
| Alphaproteobacteria | | | | 1 | | | 1 | | 3 | | | | | | | | 1 | | | | | | |
| Betaproteobacteria | | | | | 1 | | | | 1 | | 1 | | | | | | 1 | | | | | | |
| Deltaproteobacteria | 3 | 22 | 3 | 1 | | 1 | | | 10 | 9 | 3 | | | 4 | | 12 | | 5 | 4 | 1 | 2 | | |
| Gammaproteobacteria | | | 2 | 1 | | | | | | | | 3 | | | | | | | | | | | |
| Epsilonproteobacteria | | 7 | | | | | | 2 | | | | 1 | | | | | | | | | | | |
| Euryarchaeota | | | | | | | | | | 2 | | | | | | | | | | | | | |
| Total | 7 | 29 | 5 | 5 | 1 | 1 | 1 | 2 | 17 | 12 | 12 | 4 | 1 | 4 | 14 | 15 | 2 | 12 | 18 | 8 | 103 | 1 | 47 |
